# Multilocus DNA barcoding – Species Identification with Multilocus Data

**DOI:** 10.1101/155861

**Authors:** Junning Liu, Jiamei Jiang, Shuli Song, Luke Tornabene, Ryan Chabarria, Gavin J P Naylor, Chenhong Li

## Abstract

Species identification using DNA sequences, known as DNA barcoding has been widely used in many applied fields. Current barcoding methods are usually based on a single mitochondrial locus, such as cytochrome c oxidase subunit I (COI). This type of barcoding is not always effective when applied to species separated by short divergence times or that contain introgressed genes from closely related species. Herein we introduce a more effective multi-locus barcoding framework that is based on gene capture and “next-generation” sequencing and provide both empirical and simulation tests of its efficacy. We examine genetic distinctness in two pairs of fishes that are sister-species: *Siniperca chuatsi* vs. *S. kneri* and *Sicydium altum* vs. *S. adelum*, where the COI barcoding approach failed species identification in both cases. Results revealed that distinctness between *S. chuatsi* and *S. kneri* increased as more independent loci were added. By contrast *S. altum* and *S. adelum* could not be distinguished even with all loci. Analyses of population structure and gene flow suggested that the two species of *Siniperca* diverged from each other a long time ago but have unidirectional gene flow, whereas the two species of *Sicydium* are not separated from each other and have high bidirectional gene flow. Simulations demonstrate that under limited gene flow (< 0.00001 per gene per generation) and enough separation time (> 100000 generation), we can correctly identify species using more than 90 loci. Finally, we selected 500 independent nuclear markers for ray-finned fishes and designed a three-step pipeline for multilocus DNA barcoding.

---

DNA barcoding has been very successfully employed in many applied fields, ranging from routine species identification (Sutou et al. 2011; Candek and Kuntner 2015; Hassold et al. 2016), to the discovery of cryptic species (Witt et al. 2006; Kadarusman et al. 2012), tracking of invasive species (Saunders 2009; Ghahramanzadeh et al. 2013; Marescaux and Van Doninck 2013), conservation, and community ecology (Tanzler et al. 2012; Nevill et al. 2013; Hartvig et al. 2015; Shapcott et al. 2015). The mitochondrial cytochrome c oxidase subunit I gene (COI) has a good amount of variation and is easy to amplify using PCR based approaches in most animal groups (Hebert et al. 2003; Smith et al. 2005; Vences et al. 2005; Ward et al. 2005). It has become the most commonly used marker for animal DNA barcoding since it was first proposed more than a decade ago (Hebert et al. 2003). In most cases, single-locus (COI) DNA barcoding results in successful species identification. For example, a success rate close to 100% were reported for Germany herpetofauna (Hawlitschek et al. 2016), more than 90% for Chinese rodents (Li et al. 2015a), more than 80% for freshwater fishes of the Congo basin (Collins et al. 2012; Decru et al. 2016), and 100% for mosquitoes (Chan et al. 2014). However, the success rate of species identification was low for species complexes with gene flow (Hawlitschek et al. 2016) or where species had only recently diverged (van Velzen et al. 2012).

In order to use barcoding for species identification, within species variation must be less than between species variation. This generates a “break” in the distribution of distances that is referred to as the “barcoding gap”. Indeed one of the common causes of barcoding failure occurs when differences in demography eliminate the barcoding gap, because intra-specific differences are greater than inter-specific differences for the clades being compared. To an extreme, two individuals could have the same COI sequence, while being distinctly different species. Shared COI haplotypes have been reported in different species of spiders (Spasojevic et al. 2016), birds (Aliabadian et al. 2013) and fishes (Mabragana et al. 2011). The single-locus barcoding is prone to misidentification when different species share haplotypes.

Although haplotypes at a single locus, such as COI can be shared between two species, it is unlikely that individuals of two species share alleles across multiple independent genes. Accordingly, multilocus data should perform better for species identification than any single locus could. Dowton et al. (2014) proposed “next-generation” DNA barcoding based on multilocus data in which they incorporated multispecies coalescent species delimitation. They analyzed *Sarcophaga* flesh flies with two loci, mitochondrial COI and nuclear carbomoylphosphate synthase (CAD), and found out that their coalescent-based *BEAST/BPP approach was more successful than standard barcoding method (Dowton et al. 2014). However, Collins and Cruickshank (2014) reanalyzed Dowton et al.’s data and showed that standard single locus (COI) barcoding method could achieve the same accuracy as the new multilocus framework did if an optimized distance threshold was applied (Brown et al. 2012; Puillandre et al. 2012; Virgilio et al. 2012; Sonet et al. 2013).

The experiment of Dowton et al. (2014) seemed unsuccessful, but the likely reason for this was that the data they used was not challenging enough for standard single-locus barcoding methods, because there was only one unidentifiable individual that was more divergent from its closest putative conspecific than the optimized threshold (Collins and Cruickshank 2014). The other reason is that only a single nuclear gene was used in the study of Dowton et al. (2014), thus providing little additional information (Collins and Cruickshank 2014).

In the past it has been challenging to obtain sequences from sufficient independent nuclear loci from a broad taxonomic group to make multilocus DNA barcoding effective, but tools for finding thousands of nuclear gene markers (Bi et al. 2012; Li et al. 2012; Hedtke et al. 2013) and collecting their sequences through cross-species gene capture and next-generation sequencing are now available (Li et al. 2013), providing an opportunity to rigorously test the power of multilocus DNA barcoding. In this work, we tested the effectiveness of multilocus barcoding employing hundreds of nuclear loci, to correctly identify species that were not distinguishable based on only COI or a few nuclear loci.

## MATERIALS AND METHODS

### Species discrimination using empirical data

We have developed 4,434 single-copy nucleotide loci for ray-finned fishes, and tested them in 83 species (29 families and 12 orders), covering major clades of ray-finned fishes. Those markers have few missing data in the taxa tested, showing promise for their deployment in phylogenetics and population genetic analyses. We adopted those 4,434 loci as candidate barcoding markers in order to further optimize a subset of universal markers for all ray-finned fishes. We choose loci that could be readily captured and sequenced across taxa, and that were variable based on their average p-distance values among taxa.

Some of the most challenging instances for DNA barcoding occur when taxa are recently diverged or when gene flow exists between closely related species, or both. In an effort to design a rigorous barcoding scheme, we picked empirical study systems that would involve both challenges. The first involved sinipercid fishes, a family of fishes containing two genera, 9 to 12 species depending on the authority referenced (Zhou et al. 1988; Li 1991; Liu and Chen 1994; Nelson 2006). Among them, two sister species, *Siniperca chuatsi* and *S. kneri* have distinct morphological characters, such as number of plyoriccaecum, ratio between eye length and head length (Zhou et al. 1988), but they are not distinguishable using mitochondrial control region sequences (Zhao et al. 2008). These two sister species are allopatric in most of their distribution regions (Zhou et al. 1988; Li 1991), so the reason for unsuccessful species identification in these sister species is likely due to their recency of speciation (Zhao et al. 2008).

The other group of fishes that we used to test the multilocus DNA barcoding method is *Sicydium. Sicydium* is a group of diadromous gobies native to fast-flowing streams and rivers of the Americas (Central America, Mexico, Cocos Island, the Caribbean, Colombia, Ecuador and Venezuela) and Africa. There are two syntopic species, *S. altum* and *S. adelum* that could be separated according to distinct dental papillae and other morphological characters (Bussing 1996), but they are indistinguishable using mitochondrial or nuclear genes (Chabarria 2015). Because these two closely related species are frequently found together (Bussing 1996), it is possible that they have been subject to interspecific gene flow which would account for the high degree of genetic similarity between them. These two pairs of sister-species were used as test cases to evaluate how gene flow and shallow divergence times might affect species discrimination and identification based on multilocus barcoding.

### Taxa sampling, target gene enrichment, sequencing and reads assembly

We collected sequence data from 16,943 loci of nine sinipercid species, including five *S. chuatsi* and five *S. kneri* (Song et al., accepted). Here, we reused this dataset for testing multilocus barcoding. Sequences of the 4,434 loci of the sinipercids were retrieved from Song et al.’s data. The samples included five *Coreoperca whiteheadi*, one *S. scherzeri*, five *S. obscura*, two *S. undulata*, three *S. roulei*, five *S. chuatsi* and five *S. kneri*.

For the goby study, nine *S. altum* and seven *S. adelum* were collected from Costa Rica. Total genomic DNA was extracted from fin clips using a Tissue DNA kit (Omega Bio-tek, Norcross, GA, USA) and the concentration of DNA was quantified using NanoDrop 3300 Fluorospectrometer (Thermo Fisher Scientific, Wilmington, DE, USA). The goby samples were enriched and sequenced for the same 4,434 loci. The amount of DNA used for library preparation was 1 μg for each sample. The DNA sample was first sheared to 250 bp using a Covaris M220 Focused-ultrasonicator^TM^ (Covaris, Inc. Massachusetts, USA). A MYbaits kit containing baits for the 4,434 loci was synthesized at MYcroarray (Ann Arbor, Michigan, USA). The baits were designed on sequences of *Oreochromis niloticus* with 3 × tiling. Blunt-end repair, adapter ligation, fill-in, pre-hybridization PCR and target gene enrichment steps followed the protocol of cross-species gene capture (Li et al. 2013). The enriched libraries were amplified with indexed primers, pooled equimolarly and sequenced on a lane of Illumina HiSeq 2500 platform with other samples. The raw reads were parsed to separate file for each species according to the indices on the adapter. Reads assembling followed the pipeline of Yuan et al. (2016). Mitochondrial COI gene of both the sinipercids and the gobies was also amplified and sequenced using Sanger sequencing to compare COI barcoding with multilocus DNA barcoding using two pairs of primers (siniF: AACCAGCGAGCATCCATCTA and siniR: CAGTGGACGAAAGCAGCAAC for the sinipercids; sicyF: GGTTGTGTTGAGGTTTCGGT and sicyR: TCCGAGCCGAACTAAGTCAA for Sicydium).

### Effect of increasing number of loci on species discrimination

Our assumption was that individuals of recently diverged species should be more discernible using many loci than using fewer loci. Thus, we calculated p-distance among 10 individuals of *Siniperca*, including five *S. chuatsi* and five *S. kneri*, using different number of loci to test this hypothesis. Loci with no missing data in all 10 individuals of *Siniperca* were picked using a custom Perl scripts (picktaxagene.pl, Supplementary Materials). The obtained 2,612 loci were then sorted by their average p-distance (distoutlier.pl, Supplementary Materials), so outlier loci with extreme large p-distance could be checked by eye to spot bad data or bad alignment. After removing the bad data, a different number (1, 3, 10, 30, 50, 70, 90, 100, 200, 300, 400, 500, 600, 700, 800, 900, 1000, 2000) of loci were randomly picked and concatenated (samplegene.pl, Supplementary Materials) for calculating p-distance among individuals (gapdis.pl, Supplementary Materials). The sampling at each level of different number of loci was repeated two hundred times. The p-distance among individuals vs. the number of loci used was drawn with GraphPad Prism 5 (San Diego, California). To check the effect of increasing number of loci on species discrimination, the “all species barcodes” criteria was applied, that is queries was considered successfully identified when they were followed by all conspecifics according to their barcode (Meier et al. 2006). Custom Perl script was used to calculate the rate of successful identification for 200 replicates at each level of number of loci used (ID_correct_rate.pl, Supplementary Materials). Among individual p-distance and rate of successful identification also were calculated for the *Sicydium*. Sequences of COI gene also were used to calculate p-distance between individuals from the same species and from different species to compare with the results of nuclear genes. Spider (Brown et al. 2012) was used to optimize barcoding distance threshold and to identify species using COI sequences as suggested by Collins and Cruickshank (2014). The final number of loci recommended for DNA barcoding was chosen based on the effect of increasing number of loci on the success rate of species discrimination.

### Estimating species divergence and gene flow in the empirical data

Gene flow and differentiation time of S. chuatsi *and* S. kneri *was estimated* using IMa2 program with 200 loci (Hey 2010). The MCMC was run for 10 million generations with sample recorded every hundred generations. The number of chains was set to 20. The running parameters were set as -q2, -m1, -t3, -b 10000000, d100, -hn20 and -s123. An additional run was performed with the same parameter but different seeds –s111. These two run showed decent mixing, and similar results, so we combined results from the two runs. Similar runs were done for the two species of *Sicydium*. The genetic differentiation between the two species of *Siniperca* and the two species of *Sicydium* also was estimated using Structure 2.3.4 (Pritchard et al. 2000). Three iterations for 100,000 generations (using a 100,000 burnin) were run for each value of K (number of population clusters) ranging from 1 to 3. To identify the number of population clusters that captures the major structure in the data, Structure Harvester (Earl and vonHoldt 2012) was used to calculate the peak value for delta K (Evanno et al. 2005).

### Simulating sister species sequence data with different divergence times and gene flow

We simulated two diverging species with various splitting time and migration rates to explore the effect of changing these two factors on species discrimination over a broader range of parameter space. According to the IMa2 results of the empirical data, the splitting time was set as 1000, 10000, 100000, and 700000 generations. The migration rate was set as 0, 0.000001, 0.00001, and 0.0001 per generation. The simulation with 1000 generations splitting time was combined with only 0 migration rate, because the two simulated species were already indistinguishable under 1000 generations splitting time even when there was no gene flow in the simulation. The simulations with 10000, 100000, and 700000 generation splitting time were combined with all four migration rates. Fastsimcoal2 (Excoffier and Foll 2011; Excoffier et al. 2013) was used to generate the simulated data. Twenty thousand replicates were simulated for each scenario. The effective population size used for simulation was 20000 in the ancestor species and the two descendant species. Five sequences were sampled from each simulated species. The simulated data were used to calculate p-distance among individuals of the same and different species. Species identification success rate applying “all species barcodes” criteria was calculated as described above. Identification success rate using different number (1, 3, 5, 10, 30, 50, 70, 90, 100, 200, 300, 400, 500, 600, 700, 800, 900, 1000) of simulated loci was plotted against species splitting time and migration rate using R (R_Core_Team 2015).

### A three-step multilocus DNA barcoding pipeline

It is straightforward to use distance based methods to reveal divergence of two sister species in the empirical and simulated data. But for more than two species, distance based species identification becomes more complicated. Firstly, an ad hoc barcoding threshold is needed to judge whether the query is one of the species represented in the database or is a new and distinct species, but sometimes no barcoding gap exists for establishing such thresholds. Secondly, the shortest distance does not guarantee a sister species relationship either, because sister species with long branches might be less similar to the query species than a non-sister species with a short branch. To avoid these risks, we propose a three-step DNA barcoding method (Fig. 1).

**Figure 1.**
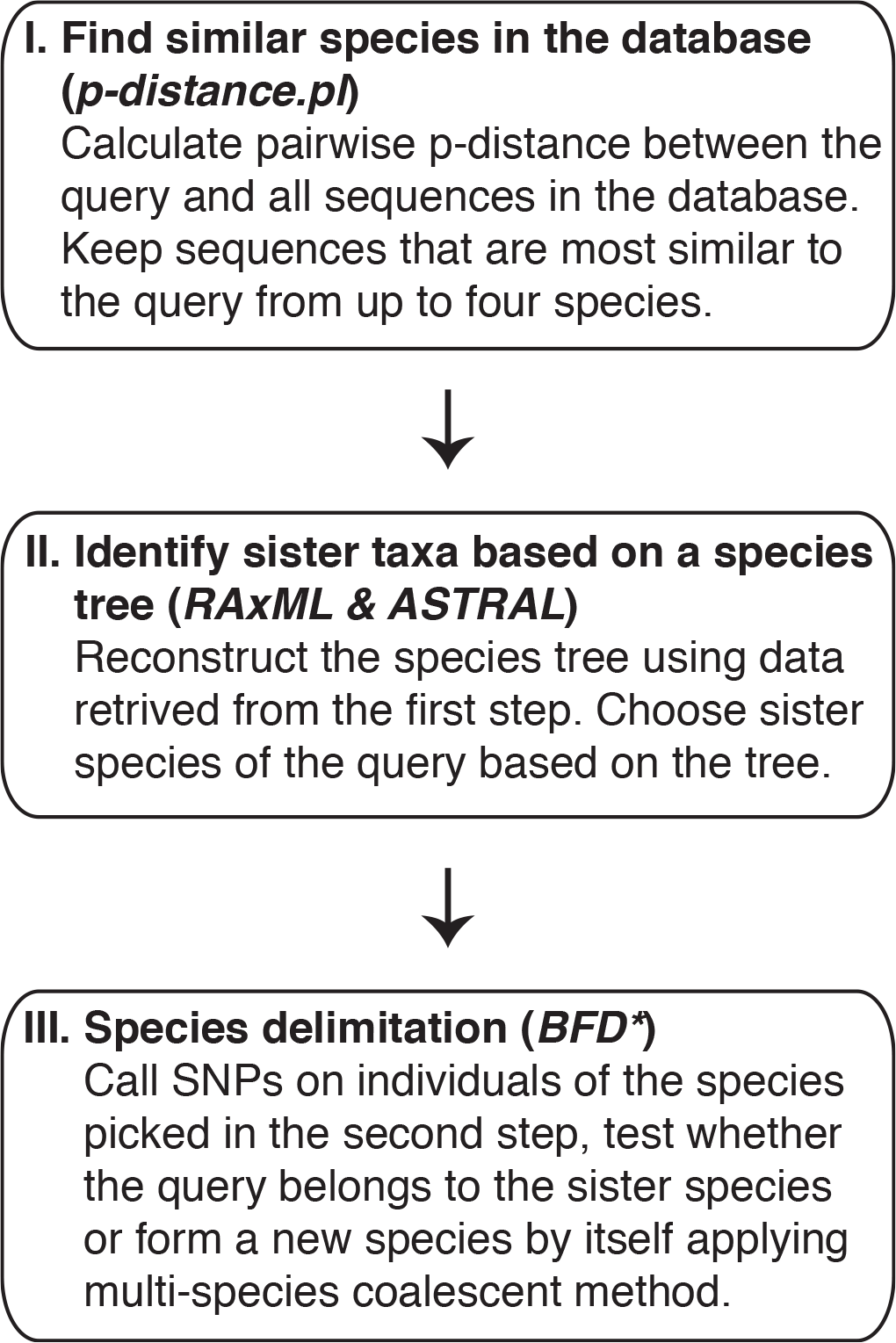
A three-step multilocus DNA barcoding pipeline.

In the first step, p-distances between the query and all sequences in the database are calculated. The sequences that are similar to the query are kept for subsequent analyses (p-distance.pl, Supplementary Materials). This is a fast screening process to retrieve all sequences from potential conspecifics or sister species. Because the closest sequence might not be from a conspecifics or sister species, sequences from up to four species are kept. In the second step, a species tree is reconstructed using the sequences from the first step to identify potential conspecifics or sister species of the query using RAxML version 8 (Stamatakis 2014) and ASTRAL 4.10.6 (Mirarab et al. 2014; Mirarab and Warnow 2015; Mirarab et al. 2016). Individual gene trees are inferred using RAxML with the GTRGAMMA model, and then a species tree is recovered from those gene trees using ASTRAL. The potential conspecifics or sister species to the query are then chosen based on the phylogenetic relationship depictured in the species tree. In the third step, species delimitation is done using a Bayes factor delimitation approach, BFD* (Leaché et al. 2014). Single nucleotides polymorphism (SNP) data are retrieved from the sequencing reads of the species chosen in step two and used for the BFD* analysis. A path sampling with 48 steps was conducted to estimate the marginal likelihood with a Markov chain Monte Carlo (MCMC) chain length of 200,000 and a pre-burnin of 50,000 following the recommended settings in BFD* (Leaché et al. 2014). The strength of support for compared hypotheses was evaluated from Bayes factor scale, 2ln(BF) using the framework of Kass and Raftery (1995). The BF scale is as follows: 0 < 2ln(BF) < 2 is not worth more than a bare mention, 2 < 2ln(BF) > 6 means positive evidence, 6 < 2ln(BF) < 10 represents strong support, and 2ln(BF) > 10 represents decisive support. If the result of BFD* analysis does not support two separate species, the query will be assigned to the “sister species”; otherwise, the query will be considered as a new species with its sequences add to the database and further study on its species status will be recommended.

The final set of selected markers was used for testing the above-described three-step multilocus DNA barcoding in the sinipercids, including 26 individuals of seven species. An individual of *S. kneri* or *S. chuatsi* was randomly chosen as unknown query that needs to be identified. The sequences of the unknown specimens and all other sequences in the database were aligned using Clustal Omega v1.1.1 (Sievers et al. 2011). Custom Perl scripts, concatnexus.pl and gapdis.pl were used to concatenate the sequences of individual loci, to calculate their p-distance between the query and the sample in the database, and sorted them by the p-distance to find all individuals that are close to the query sample.

### Testing effect of missing data in the database or in the query on the success rate of species identification

To test if our method could identify new species when the sequences of conspecifics are not in the database, all samples of *S. kneri* were removed from the database except that one random selected *S. kneri* individual was left as query. To access the effect of missing data in the query sample, one *S. kneri* was selected as an unknown sample, and 20 percent, 30 percent, and 50 percent of its loci were excluded, then the data were used for multilocus DNA barcoding analysis.

## RESULTS

We investigated effect of increasing number of loci on species discrimination and identification using empirical data (between *Siniperca chuatsi* and *S. kneri* and between *Sicydium altum* and *S. adelum)*. We subsequently estimated the population parameters, gene flow and divergence time for both pairs of species. Guided by the patterns seen in the empirical data, we simulated sequences with different splitting times and migration rates, and explored the effect of divergence time and gene flow on the success rate of species identification over a broader range of the relevant parameter space. Finally, we selected 500 nuclear markers for ray-finned fishes and designed a three-step pipeline for multilocus DNA barcoding.

### Species discrimination using empirical data

After all loci with missing taxa were excluded, 2586 loci were retained for *Siniperca*. The intra- and interspecific p-distances between five individuals of *S. chuatsi* and five *S. kneri* using different numbers of nuclear loci or COI are shown in Figure 2. The intraspecific p-distance (red) calculated using one locus or a small number of loci overlap with interspecific p-distance (blue). There is no barcoding gap separating the intra- and interspecific distances. Intraspecific distances did not become distinguishable from interspecific distances until more than 90 loci were used. The gap separating the intra- and interspecific distance increased as more loci were added, but had little effect after 400 loci were used. The variance of the intra- and interspecific p-distance decreased when more loci were included in calculating the p-distance. The p-distance calculated on COI sequences also had mixed intra- and interspecific values but they were an order of magnitude higher than those calculated using nuclear loci (Fig. 2 right y-axis).

**Figure 2.**
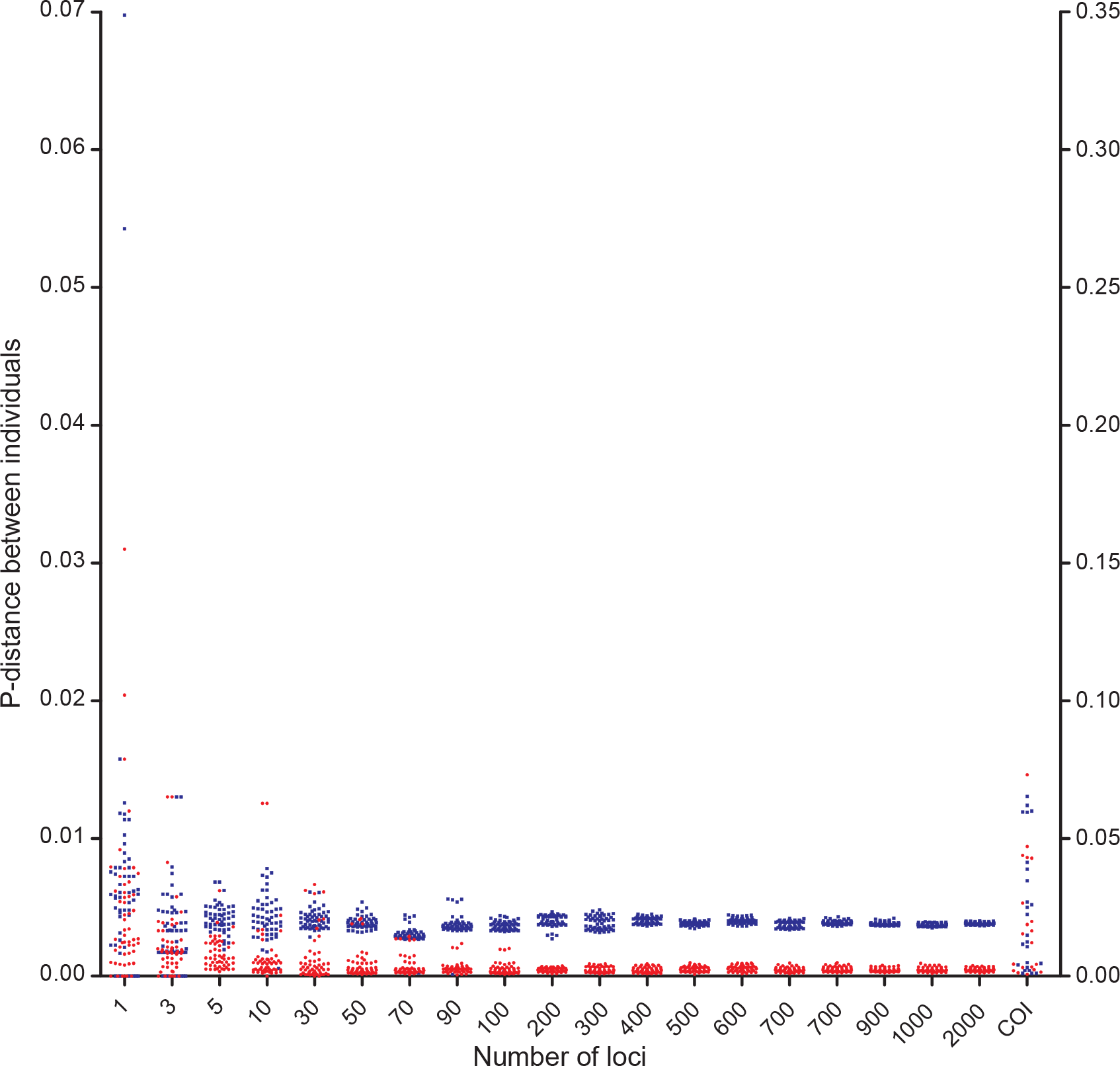
Intra- (red) and interspecific (blue) p-distance of *Siniperca chuatsi* and *S. kneri* calculated using different number of nuclear loci or COI gene. The distances based on COI (right y-axis) are one order of magnitude higher than distances calculated using nuclear loci (left y-axis).

Similar p-distance calculations on *S. altum* and *S. adelum* resulted in a different pattern than the one observed for *Siniperca*. The Intra- (red) and interspecific (blue) p-distances in *Sicydium* were always mixed together, no matter how many loci were included in the analysis. The variance of intra- and interspecific p-distance decreased when more loci were included. The intra- and interspecific p-distances calculated using COI also were indistinguishable (Fig. 3).

**Figure 3.**
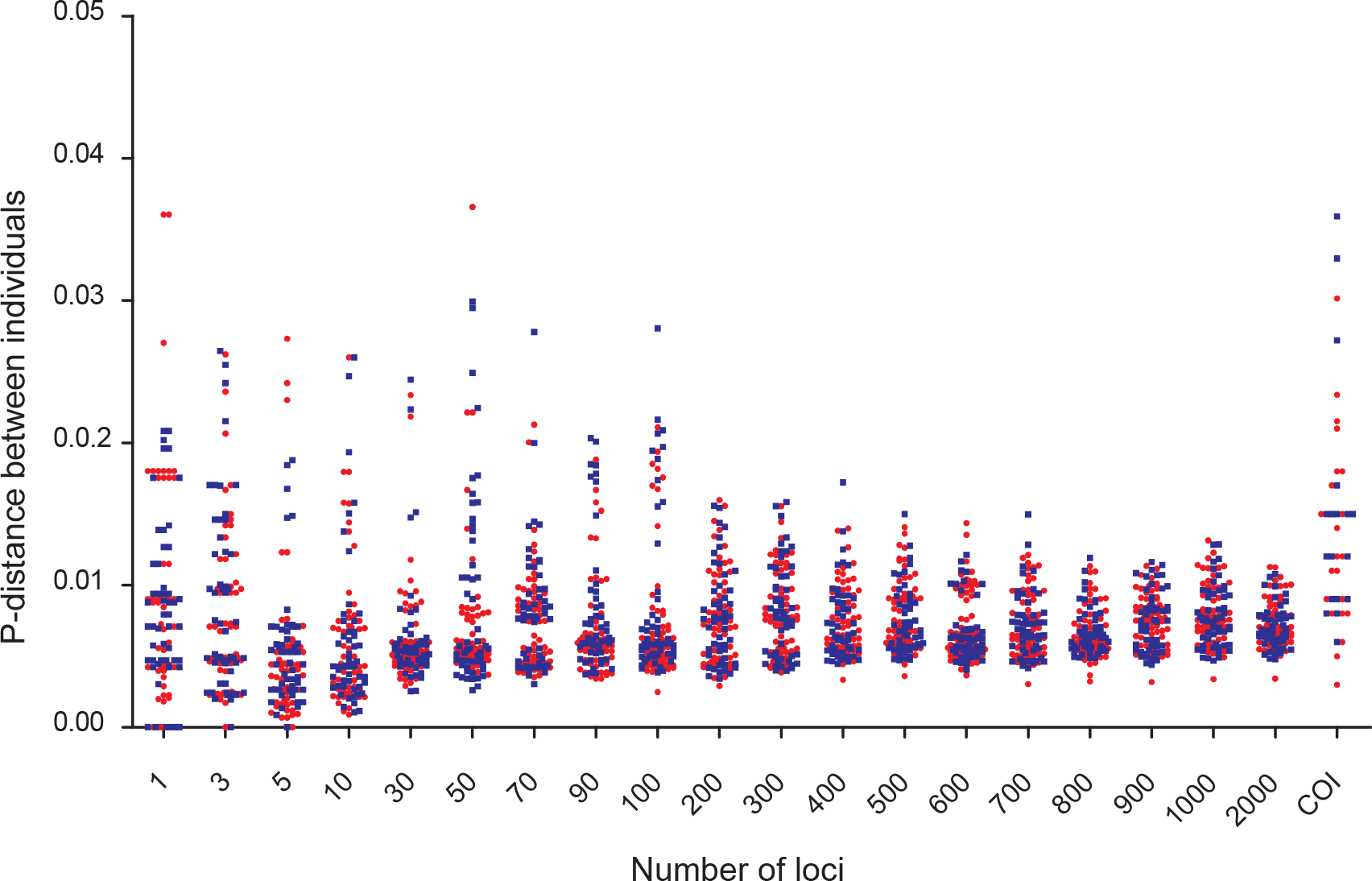
Intra- (red) and interspecific (blue) p-distance of *Sicydium altum* and *S. adelum* calculated using different numbers of nuclear loci or COI gene.

The success rate of identification was low (0.412) in *Siniperca* when only one locus was used based on “all species barcodes” criterion with two hundred trials, but it rose up quickly and reached 1.0 after more than 90 loci were added to the dataset (green dots, Fig. 4; Table S1). The identification success rate was zero in *Sicydium* according to the “all species barcodes” criterion, no matter how many loci were included in the analysis (red triangles, Fig. 4). We also applied the COI barcoding approach with an optimized threshold. The success rate of species identification using COI was zero in both *Siniperca* and *Sicydium*.

**Figure 4.**
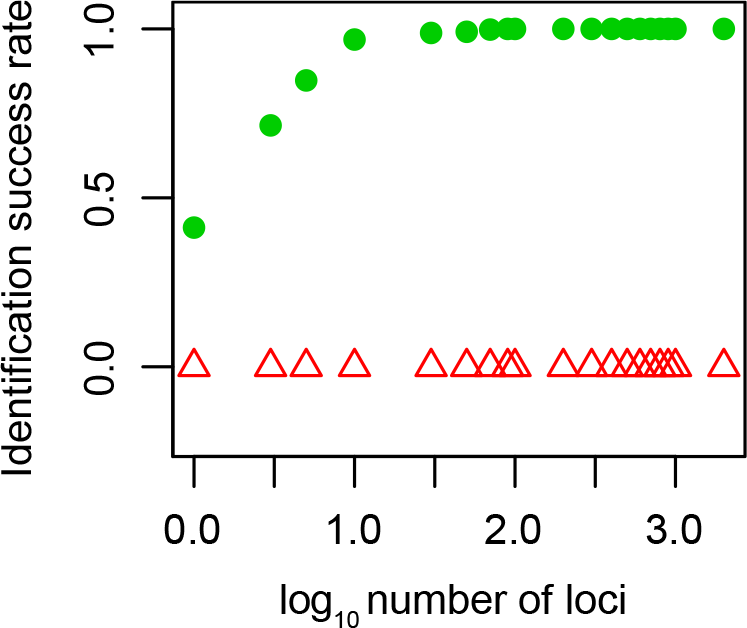
The relationship between number of loci used and success rate of identification between *Siniperca chuatsi* and *S. kneri* (green dots), and between *Sicydium altum* and *S. adelum* (red triangles).

### Population parameters inferred for the two species pairs

To investigate the difference seen in the results of the *Siniperca* and *Sicydium* analyses, we explored some of the population attributes associated with each of these two groups. Structure analysis showed that K equaled to 2 had the highest probability when analyzing the two *Siniperca* species (Fig. S1 a), but the two *Sicydium* species were indistinguishable (Fig. S1 b). The divergence time between *S. chuatsi* and *S. kneri* was estimated as t_0_ = 1.754, which would be equal to ~800,000 generations if we assume an average locus size of 300 bp, a generation time of 2 - 3 years for *Siniperca* and a substitution rate of 2.22 × 10^-9^ per site per year (Kumar and Subramanian 2002). Gene flow from *S. chuatsi* to *S. kneri* was 0.157 (not significant by LLRtest), but gene flow from *S. kneri* to *S. chuatsi* was highly significant, 0.640 (p < 0.001). The divergence time between *S. altum* and *S. adelum* was estimated as t_0_ = 0.003195, which was not significantly different from zero (HPD95_Lo_=0). Gene flow from *S. altum* to *S. adelum* was 0.494, and gene flow from *S. adelum* to *S. altum* was 0.502.

### Simulation results

To explore the effect of divergence time and gene flow on the success rate of species identification, we conducted a series of simulations using twenty thousand loci for two species with a range of splitting times and migration profiles. Five sequences from each species were sampled to calculate species identification success rate. Different number of simulated loci were randomly picked and used to identify species. The identification success rate rose with increasing number of loci included in the analyses in all scenarios (Fig. 5). When there was no migration between the two simulated species, the identification success rate increased with splitting time (Fig. 5a). The simulation with a splitting time of 1000 generations had the worst identification success rate, only 0.111 even with 1000 loci used (green circle, Fig. 5 a; Table S2). The samples with a splitting time of 10000 generations had low success rates with a small number of loci used, but rose to 1 when more than 400 loci were added to the analyses (blue triangles, Fig. 5 a). The samples with a splitting time of 100000 generations had a success rate of 1 when more than 10 loci were used (black crosses, Fig. 5a). Samples with a splitting time of 700000 had success rate of 1 for all analyses (red line, Fig. 5a).

**Figure 5.**
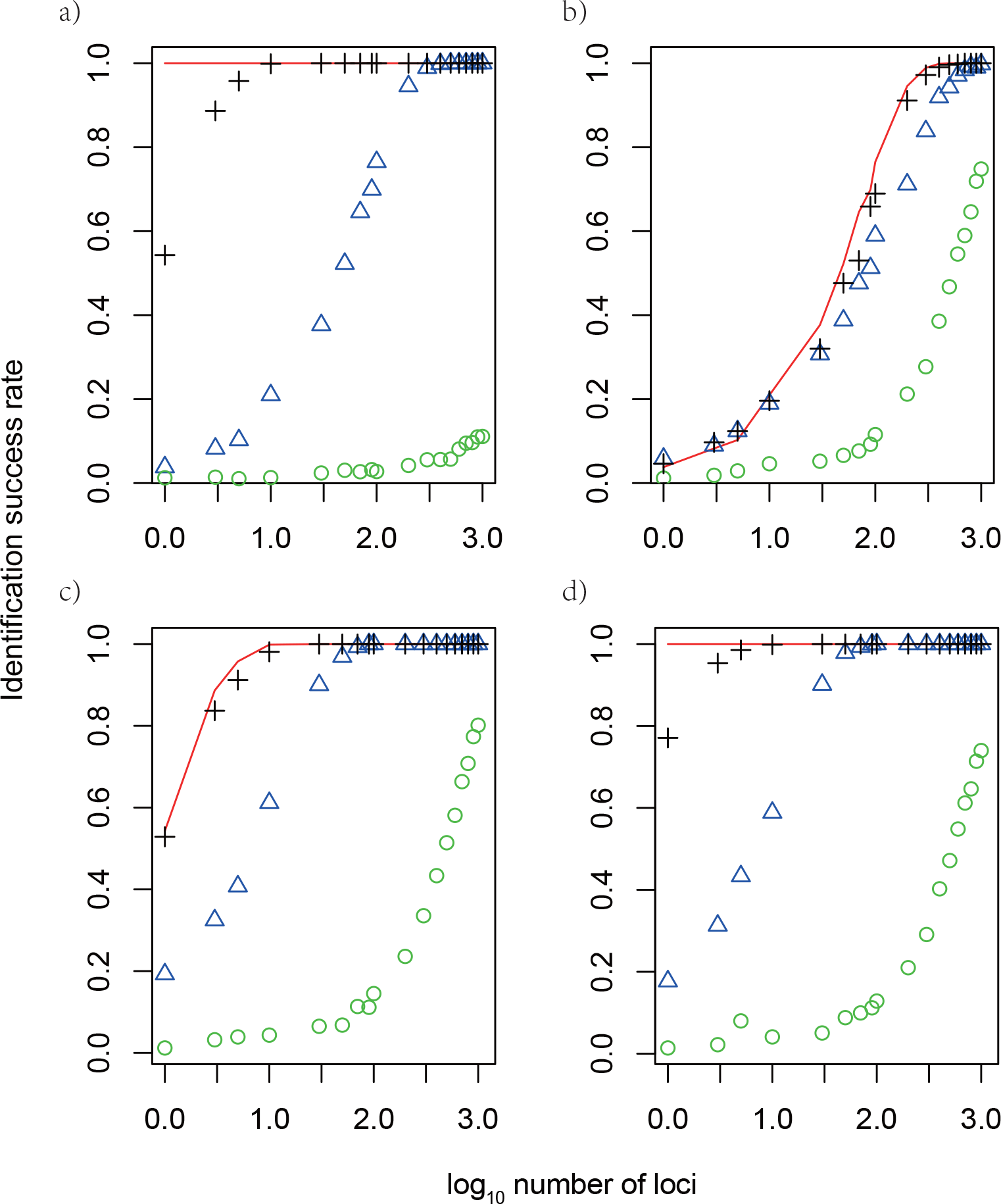
Identification success rate using simulated sequences under different scenarios. a) migration rate equals zero and divergence time equals 700000 generations (red line), 100000 generations (black crosses), 10000 generations (blue triangles), and 1000 generations (green circles); b) divergence time equals 10000 and migration rate equals 0 (red line), 0.000001 (black crosses), 0.00001 (blue triangles) and 0.0001 (green circles); c) divergence time equals 100000 and migration rate equals 0 (red line), 0.000001 (black crosses), 0.00001 (blue triangles) and 0.0001 (green circles); d) divergence time equals 700000 and migration rate equals 0 (red line), 0.000001 (black crosses), 0.00001 (blue triangles) and 0.0001 (green circles).

When gene flow was considered, high gene flow worked in concert with shallow divergence time to reduce the identification success rate (Fig. 5 b-d). A migration rate of 0.0001 (per gene per generation) always led to the worst success rate, and failed to reach a success rate of 1.0 even when all 1000 loci were used in analyses (green circles, Fig. 5 b-d; Table S3-S5). At a migration rate of 0.00001 (blue triangles, Fig. 5 b-d) or a migration rate of 0.000001 (black crosses, Fig. 5 b-d), the identification success rate improved quickly with increasing number of loci (Fig. 5 b-d; Table S3-S5). When the divergence time was greater than 100000 generations and gene flow was lower than 0.00001, the identification success rate reached 1.0 when more than 90 loci were added to the analysis (Fig. 5 c-d; Table S4-S5).

To test whether the length of sequence or the number of loci was the key for success in species identification, we simulated a single locus with increasing size matching the total length of multiple loci. We found that increasing the length of a single locus from 300 bp to 9000 bp improved the success rate slightly, but the success rate did not change when longer sequences were used (Fig. 6 red circles; Table S6). In contrast, concatenating more independent loci with the same total length as the single locus continuously improved the identification success rate, until it reached one (Fig. 6 blue triangles; Table S6).

**Figure 6.**
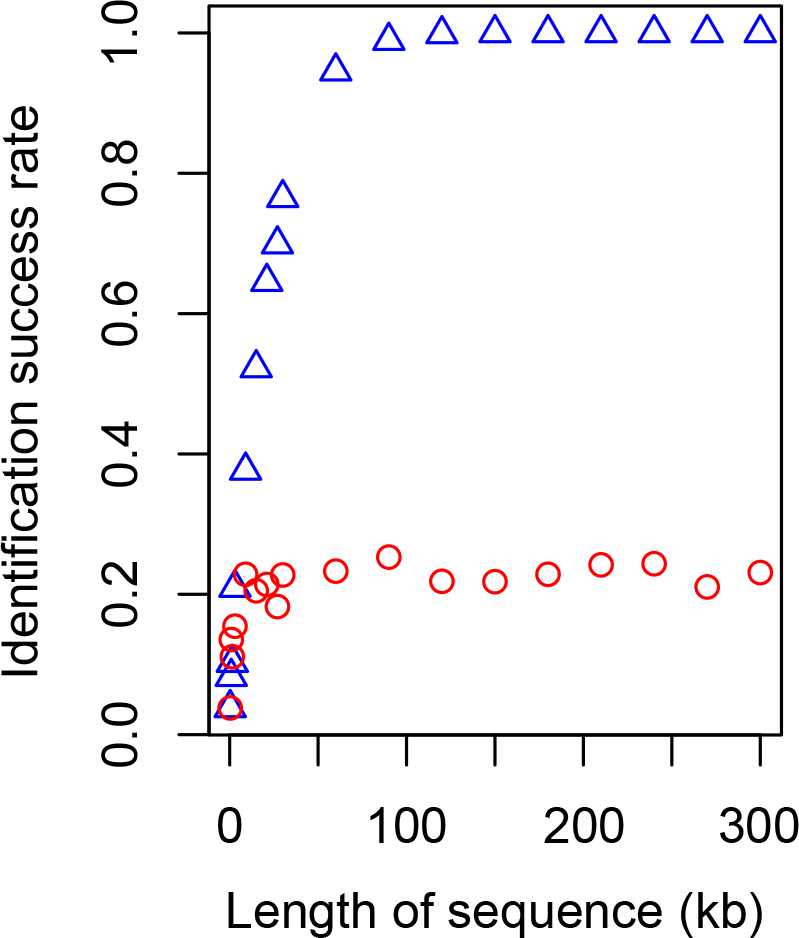
Comparison between success rates of species identification based on a) a single locus and b) multiple loci. The length of the single locus equals the total length of multiple loci (300 bp each).

### Multilocus DNA barcoding using empirical data

Based on the results from p-distance and species identification analyses of simulated and empirical data, we decided to pick 500 loci for multilocus DNA barcoding. First, we filtered the 4,434 markers developed for all ray-finned fishes and kept 750 loci with the lowest number of missing taxa. Next, we sorted the 750 loci by their average p-distance and picked from them 500 independent loci with large p-distances. This design was implemented both to minimize missing data when applying to ray-finned fishes and to ensure that loci would be variable enough for multilocus DNA barcoding. Information describing the 500 loci is listed in Supplementary Materials (Table S7).

Three individuals, 839_3 *(S. kneri)*, 839_6 *(S. kneri)*, and 938_1 *(S. chuatsi)* were randomly selected. Each of the randomly picked individuals was used to simulate “a putatively unknown” query for identification. Firstly, the p-distance between the unknown query and the other sinipercids in the database was calculated (Table S8). Secondly, based on the sorted list of p-distances, we selected five closely related taxa, including the query. For example, for 839_3, we used sequence data of 839_3, *S. kneri, S. chuatsi, S. undulata* and *S. obscura* to reconstruct a species tree, in which 839_3 was found to be sister to *S. kneri* (Fig. S2). We then ran a BFD* test to delimitate the unknown query (839_3) with *S. kneri* using *S. chuatsi* as outgroup. The BFD* analyses correctly grouped 839_3 (S. *kneri)* with *S. kneri* (Table 1). The two other randomly picked samples, 839_6 (S. *kneri)*, and 938_1 (S. *chuatsi)* were also correctly identified (Table S8 and S9).

**TABLE 1.** Results for species delimitation on unknown sample 893_3 *(S. kneri)* using BFD* based on all 500 nuclear loci, missing 20%, 30% and 50% of the 500 loci or missing conspecific of *S. kneri* in the database.

| Data treatment | Model | Marginal likelihood | 2lnBF |
| --- | --- | --- | --- |
| Using all data | Lumping 839_3 and *S. kneri* | -1575.80 | 20.62 |
|  | Splitting 839_3 and *S. kneri* | -1586.11 |  |
| Excluding conspecifics of 839_3 | Lumping 839_3 and *S. chuatsi* | -2350.77 |  |
|  | Splitting 839_3 and *S. chuatsi* | -1222.90 | 2255.7 |
| Excluding 20% loci of the 839_3 | Lumping 839_3 and *S. kneri* | -1467.41 | 26.54 |
|  | Splitting 839_3 and *S. kneri* | -1480.68 |  |
| Excluding 30% loci of the 839_3 | Lumping 839_3 and *S. kneri* | -1247.44 | 22.12 |
|  | Splitting 839_3 and *S. kneri* | -1258.50 |  |
| Excluding 50% loci of the 839_3 | Lumping 839_3 and *S. kneri* | -914.60 | 22.40 |
|  | Splitting 839_3 and *S. kneri* | -925.80 |  |

DNA barcoding using only COI data was unsuccessful. In many cases, the closest taxa of the unknow samples were not their conspecifics either in the tree or measured by p-distances (Fig. S3, Table S10).

### Effect of missing data on multilocus DNA barcoding

When all conspecifics were excluded from the database, the unknown query, 893_3 (S. *kneri)* was found to be closely related to its sister species *S. chuatsi* (Fig. S4). The p-distances also indicated that the unknown was related to *S. chuatsi* (Table S11). A species delimitation analysis was run with the BFD* method to test whether the unknown should be assigned to *S. chuatsi* or not. The result of BFD* strongly support the unknown query as a separate species (2lnBF = 2255.7; Table 1). In other tests, we keep the database intact, but excluded 20%, 30% and 50% of the loci from the unknown query (893_3 *S. kneri)*. We still identified the unknown correctly using the multilocus DNA barcoding approach (Table 1).

## DISCUSSION

### Species are more distinguishable with more independent loci than with a few

Our results demonstrated that the difference between species become more distinct when more independent loci are used. The Intra- (red) and interspecific (blue) p-distance between individuals of *S. chuatsi* and *S. kneri* were largely overlapping when only COI gene or a few randomly picked nuclear gene were used to calculate the p-distance (Fig. 2). When more loci were added to the analyses, the intra- and interspecific distance became better separated. At 90 loci, a “barcoding gap” between the intra- and interspecific distance emerged. The variance of the intra- and interspecific distances also decreased as the number of loci used in the analyses increased. Based on these findings we conclude that the lack of an apparent barcoding gap between *S. chuatsi* and *S. kneri* using COI or a few nuclear genes is due to sampling error. Using more independent loci would likely improve the estimates of population parameters (Lee and Edwards 2008). Similarly, more indepenent loci should improve precision of both the estimated intra- and interspecific genetic distance, resulting in increased discriminatory power (Fig. 4). The same patterns were obeserved in all of our simulated analyses, namely that the species identification success rate rose with increasing number of loci (Fig. 5). Interestingly, using longer genes instead of more genes did not improve species identification (Fig. 6).

### Divergence time vs. gene flow

Gene flow between sister species can cause problems that are similar to those caused by a lack of divergence. *S. chuatsi* and *S. kneri* were estimated split at around 80 thousand generations ago, with uni-directional introgression flowing from *S. kneri* to *S. chuatsi*, m_1>0_ = 0.640. Therefore, the lack of reciprocal monophyly or barcoding gap between *S. chuatsi* and *S. kneri* using COI or a few nuclear loci could, in fact, be caused by gene flow between the two species rather than the short divergence time originally hypothesized.

*Sicydium altum* and *S. adelum* were estimated to have split very recently, t_0_ = 0.003195. Bi-directional gene flow was estimated as 0.494 from *S. altum* to *S. adelum*, and 0.502 from *S. adelum* to *S. altum*. All of our analyses could not differentiate between *S. altum* and *S. adelum* genetically. Structure analysis (Fig. S1), species identification and p-distance assessments (Fig. 3 and 4) all indicated that *S. altum* and *S. adelum* are indistinguishable. Accordingly, we suggest that the taxonomic status of *S. altum* and *S. adelum* be revisited by a more detailed morphological analysis.

It is difficult to tell whether gene flow or short species divergence time played a more prominent role in obstructing DNA barcoding. It has been reported that a considerable proportion of animal species do not form monophyletic groups (Funk and Omland 2003; Ross 2014), but the causes for such patterns have not yet been fully explored. From the results of our empirical and simulated analyses, we conclude that when the splitting time between sister species was more than 100000 generations old and the migration rate was lower than 0.00001, using multilocus DNA barcoding (with more than 90 loci) we could correctly determine the species status of unknown samples, whereas single-locus DNA barcoding suffered from lacking of power in species discrimination.

### Markers for multilocus DNA barcoding

A suite of universal gene markers that could be used on a whole group of organisms is a prerequisite for multilocus DNA barcoding. Because of improvements in sequencing technology and the increasing number of publicly accessable genome data bases, more and more genome-scale markers have been developed for different group of organisms, such as turtles (Shen et al. 2011), birds (McCormack et al. 2013), tapeworms (Yuan et al. 2016), flower flies (Young et al. 2016), plants (Schmickl et al. 2016), echinoderms (Hugall et al. 2016), insects (Blaimer et al. 2016) and vertebrates (Li et al. 2013). Some of these markers can be applied across broad groups of organisms, whereas other have only been tested for restricted groups. We predict that obtaining suitable sets of markers for multilocus DNA barcoding will not be a limitation, but a lot of testing will need to carried out across a broad range of taxa before an agreed set of common markers can be established for each major group of organisms.

Our pick of 500 markers for ray-finned fishes has been tested in all major lineages of fishes. We chose markers that were found to be present in most groups of fishes and that were variable across groups. We recommend using them as standard multilocus DNA barcode markers for all ray-finned fishes. Our results indicate that more than 90 loci should be enough for species identification, but we advocate using the complete set of 500 loci, as there is almost no extra cost in capturing 500 rather than 90 loci. Additionally, targeting more loci provides insurance against missing data. We found that missing 20%, 30% and up to 50% loci in the unknown sample had no effect in identification success.

Other alternatives to collecting large datasets for DNA barcoding include genome skimming (Coissac et al. 2016) and whole-chloroplast genome sequencing (Li et al. 2015b). Genome skimming employs low-coverage shotgun sequencing of genomic DNA, which circumvents the need for PCR, avoiding the needs for univerals primers. Because genome skimming is unselective, it involves collecting a lot of data that ultimately is not used, but requires data storage and analysis resources. Low-coverage shotgun sequencing also yields a high proportion of missing data. Sequencing genomes of chloroplasts or other organelles is focused on a single long sequence, which tends to yield low success rate of species identification, as shown in our simulation.

### Three-step data analysis pipeline of multilocus barcoding

Dowton et al. (2014) proposed an pipeline integrating species tree reconstruction and species delimitation. They used Beast* to build a species tree (Heled and Drummond 2010), and took the species tree as guide tree for delimitating species using BPP (Rannala and Yang 2003; Yang and Rannala 2010). Our method is similar to the method of Dowton et al. (2014). We first screened the reference database for individuals from closely related speices based on p-distance between the unknow query and sequences in the database. We only choose four closely related species as potential conspecies or sister species. We think the number of species selected is enough for the current study, because our p-distance calculation was based on many independent loci, which reduces random error. The small number of selected species could also help to relieve the computational burden associated with reconstructing the species tree in the second step. Using a combination of RAxML and ASTRAL program, we could reconstruct a specis tree of five taxa, four selected species plus the query in minutes using 500 loci. In the last step, we included only three taxa, one conspecific or sister species, one outgroup species and the unknown query for species delimiation using BFD*, which also saved computation time. We anticipate that the computational burden associated with multilocus DNA barcoding will be further reduced as new algorithms are developed, to make multilocus barcoding a real-time tool.

### Cost of multilocus DNA barcoding

Finally, from a practical standpoint, multilocus barcoding through target gene enrichment is efficient. We estimate around $90 for the total cost of capturing and sequencing 500 loci per sample, which is less than the cost of amplifying and sequencing 10 loci using the tranditional methods of PCR and Sanger sequening. The cost of target gene capture comprises: library prep, $50; RNA baits, $32; and sequencing, $8 per sample. The major costs are associated with the purchase of commercial RNA bait kits and the library preparation step, which can be lowered by purchasing kits in bulk and by using robots to automate library preparation.

## SUPPLEMENTARY MATERIAL

Data available from the Dryad Digital Repository: http://dx.doi.org/XXX.

## Acknowledgements

### ACKNOWLEDGEMENTS

The authors would like to thank Shanghai Oceanus Supercomputing Center (SOSC) for providing computational resources.

## References

Aliabadian M, Beentjes KK, Roselaar CS, van Brandwijk H, Nijman V, Vonk R. 2013. DNA barcoding of Dutch birds. Zookeys:25–48.

Bi K, Vanderpool D, Singhal S, Linderoth T, Moritz C, Good JM. 2012. Transcriptome-based exon capture enables highly cost-effective comparative genomic data collection at moderate evolutionary scales. BMC Genomics, 13:403.

Blaimer BB, Lloyd MW, Guillory WX, Brady SG. 2016. Sequence Capture and Phylogenetic Utility of Genomic Ultraconserved Elements Obtained from Pinned Insect Specimens. PLoS One, 11:e0161531.

Brown SD, Collins RA, Boyer S, Lefort MC, Malumbres-Olarte J, Vink CJ, Cruickshank RH. 2012. Spider: an R package for the analysis of species identity and evolution, with particular reference to DNA barcoding. Mol Ecol Resour, 12:562–565.

Bussing WA. 1996. Sicydium adelum, a new species of gobiid fish (Pisces: Gobiidae) from Atlantic slope streams of Costa Rica. Rev. Biol. Trop., 44:819–825.

Candek K, Kuntner M. 2015. DNA barcoding gap: reliable species identification over morphological and geographical scales. Mol Ecol Resour, 15:268–277.

Chabarria RE. 2015. Evolution of the genus Sicydium (Gobiidae: Sicydiinae).Department of Life Sciences. Corpus Christi, TX, Texas A&M University - Corpus Christi.

Chan A, Chiang LP, Hapuarachchi HC, Tan CH, Pang SC, Lee R, Lee KS, Ng LC, Lam-Phua SG. 2014. DNA barcoding: complementing morphological identification of mosquito species in Singapore. Parasite Vector, 7:569.

Coissac E, Hollingsworth PM, Lavergne S, Taberlet P. 2016. From barcodes to genomes: extending the concept of DNA barcoding. Molecular ecology, 25:1423–1428.

Collins RA, Armstrong KF, Meier R, Yi Y, Brown SD, Cruickshank RH, Keeling S, Johnston C. 2012. Barcoding and border biosecurity: identifying cyprinid fishes in the aquarium trade. PLoS One, 7:e28381.

Collins RA, Cruickshank RH. 2014. Known knowns, known unknowns, unknown unknowns and unknown knowns in DNA barcoding: a comment on Dowton et al. Syst. Biol., 63:1005–1009.

Decru E, Moelants T, De Gelas K, Vreven E, Verheyen E, Snoeks J. 2016. Taxonomic challenges in freshwater fishes: a mismatch between morphology and DNA barcoding in fish of the north-eastern part of the Congo basin. Mol Ecol Resour, 16:342–352.

Dowton M, Meiklejohn K, Cameron SL, Wallman J. 2014. A preliminary framework for DNA barcoding, incorporating the multispecies coalescent. Syst. Biol., 63:639–644.

Earl DA, vonHoldt BM. 2012. STRUCTURE HARVESTER: a website and program for visualizing STRUCTURE output and implementing the Evanno method. Conservation Genetics Resources, 4:359–361.

Evanno G, Regnaut S, Goudet J. 2005. Detecting the number of clusters of individuals using the software STRUCTURE: a simulation study. Molecular ecology, 14:2611–2620.

Excoffier L, Dupanloup I, Huerta-Sanchez E, Sousa VC, Foll M. 2013. Robust demographic inference from genomic and SNP data. PLoS Genet, 9:e1003905.

Excoffier L, Foll M. 2011. fastsimcoal: a continuous-time coalescent simulator of genomic diversity under arbitrarily complex evolutionary scenarios. Bioinformatics, 27:1332–1334.

Funk DJ, Omland KE. 2003. SPECIES-LEVEL PARAPHYLY AND POLYPHYLY: Frequency, Causes, and Consequences, with Insights from Animal Mitochondrial DNA. Annu. Rev. Ecol. Evol. Syst., 34:397–423.

Ghahramanzadeh R, Esselink G, Kodde LP, Duistermaat H, van Valkenburg JL, Marashi SH, Smulders MJ, van de Wiel CC. 2013. Efficient distinction of invasive aquatic plant species from non-invasive related species using DNA barcoding. Mol Ecol Resour, 13:21–31.

Hartvig I, Czako M, Kjaer ED, Nielsen LR, Theilade I. 2015. The Use of DNA Barcoding in Identification and Conservation of Rosewood (Dalbergia spp.). PLoS One, 10:e0138231.

Hassold S, Lowry PP, 2nd, Bauert MR, Razafintsalama A, Ramamonjisoa L, Widmer A. 2016. DNA Barcoding of Malagasy Rosewoods: Towards a Molecular Identification of CITES-Listed Dalbergia Species. PLoS One, 11:e0157881.

Hawlitschek O, Moriniere J, Dunz A, Franzen M, Rodder D, Glaw F, Haszprunar G. 2016. Comprehensive DNA barcoding of the herpetofauna of Germany. Mol Ecol Resour, 16:242–253.

Hebert PD, Cywinska A, Ball SL, deWaard JR. 2003. Biological identifications through DNA barcodes. Proc Biol Sci, 270:313–321.

Hedtke SM, Morgan MJ, Cannatella DC, Hillis DM. 2013. Targeted enrichment: maximizing orthologous gene comparisons across deep evolutionary time. PLoS One, 8:e67908.

Heled J, Drummond AJ. 2010. Bayesian inference of species trees from multilocus data. Mol Biol Evol, 27:570–580.

Hey J. 2010. Isolation with migration models for more than two populations. Mol Biol Evol, 27:905–920.

Hugall AF, O'Hara TD, Hunjan S, Nilsen R, Moussalli A. 2016. An Exon-Capture System for the Entire Class Ophiuroidea. Mol Biol Evol, 33:281–294.

Kadarusman, Hubert N, Hadiaty RK, Sudarto, Paradis E, Pouyaud L. 2012. Cryptic diversity in Indo-Australian rainbowfishes revealed by DNA barcoding: implications for conservation in a biodiversity hotspot candidate. PLoS One, 7:e40627.

Kass RE, Raftery AE. 1995. Bayes factors. Journal of the American Statistical Association 90:773–795.

Kumar S, Subramanian S. 2002. Mutation rates in mammalian genomes. Proceedings of the National Academy of Sciences of the United States of America, 99:803–808.

Leaché AD, Fujita MK, Minin VN, Bouckaert RR. 2014. Species delimitation using genome-wide SNP data. Syst Biol, 63:534–542.

Lee JY, Edwards SV. 2008. Divergence across Australia's Carpentarian barrier: statistical phylogeography of the red-backed fairy wren (Malurus melanocephalus). Evolution, 62:3117–3134.

Li C, Hofreiter M, Straube N, Corrigan S, Naylor GJ. 2013. Capturing protein-coding genes across highly divergent species. Biotechniques, 54:321–326.

Li C, Riethoven JJ, Naylor GJ. 2012. EvolMarkers: a database for mining exon and intron markers for evolution, ecology and conservation studies. Mol Ecol Resour, 12:967–971.

Li J, Zheng X, Cai Y, Zhang X, Yang M, Yue B. 2015a. DNA barcoding of Murinae (Rodentia: Muridae) and Arvicolinae (Rodentia: Cricetidae) distributed in China. Mol Ecol Resour, 15:153–167.

Li S. 1991. Geographical distribution of the Sinipercine fishes. Chinese Journal of Zoology, 26:40–44.

Li X, Yang Y, Henry RJ, Rossetto M, Wang Y, Chen S. 2015b. Plant DNA barcoding: from gene to genome. Biol Rev Camb Philos Soc, 90:157–166.

Liu H, Chen Y. 1994. Phylogeny of the sinipercine fishes with some taxonomic notes. Zoological Research, 15:1–12.

Mabragana E, Diaz de Astarloa JM, Hanner R, Zhang J, Gonzalez Castro M. 2011. DNA barcoding identifies Argentine fishes from marine and brackish waters. PLoS One, 6:e28655.

Marescaux J, Van Doninck K. 2013. Using DNA barcoding to differentiate invasive Dreissena species (Mollusca, Bivalvia). Zookeys:235–244.

McCormack JE, Harvey MG, Faircloth BC, Crawford NG, Glenn TC, Brumfield RT. 2013. A phylogeny of birds based on over 1,500 loci collected by target enrichment and high-throughput sequencing. PLoS One, 8:e54848.

Meier R, Shiyang K, Vaidya G, Ng PK. 2006. DNA barcoding and taxonomy in Diptera: a tale of high intraspecific variability and low identification success. Syst. Biol., 55:715–728.

Mirarab S, Bayzid MS, Warnow T. 2016. Evaluating Summary Methods for Multilocus Species Tree Estimation in the Presence of Incomplete Lineage Sorting. Syst. Biol., 65:366–380.

Mirarab S, Reaz R, Bayzid MS, Zimmermann T, Swenson MS, Warnow T. 2014. ASTRAL: genome-scale coalescent-based species tree estimation. Bioinformatics, 30:i5411–548.

Mirarab S, Warnow T. 2015. ASTRAL-II: coalescent-based species tree estimation with many hundreds of taxa and thousands of genes. Bioinformatics, 31:i44–52.

Nelson JS. 2006. Fishes of the world. 4th ed. New York, John Wiley and Sons, Inc.

Nevill PG, Wallace MJ, Miller JT, Krauss SL. 2013. DNA barcoding for conservation, seed banking and ecological restoration of Acacia in the Midwest of Western Australia. Mol Ecol Resour, 13:1033–1042.

Pritchard JK, Stephens M, Donnelly P. 2000. Inference of population structure using multilocus genotype data. Genetics, 155:945–959.

Puillandre N, Lambert A, Brouillet S, Achaz G. 2012. ABGD, Automatic Barcode Gap Discovery for primary species delimitation. Molecular ecology, 21:1864–1877.

R_Core_Team. 2015. R: A language and environment for statistical computing. Vienna, Austria., R Foundation for Statistical Computing.

Rannala B, Yang Z. 2003. Bayes estimation of species divergence times and ancestral population sizes using DNA sequences from multiple loci. Genetics, 164:1645–1656.

Ross HA. 2014. The incidence of species-level paraphyly in animals: a re-assessment. Mol. Phylogenet. Evol., 76:10–17.

Saunders GW. 2009. Routine DNA barcoding of Canadian Gracilariales (Rhodophyta) reveals the invasive species Gracilaria vermiculophylla in British Columbia. Mol Ecol Resour, 9 Suppl s1:140–150.

Schmickl R, Liston A, Zeisek V, Oberlander K, Weitemier K, Straub SC, Cronn RC, Dreyer LL, Suda J. 2016. Phylogenetic marker development for target enrichment from transcriptome and genome skim data: the pipeline and its application in southern African Oxalis (Oxalidaceae). Mol Ecol Resour, 16:1124–1135.

Shapcott A, Forster PI, Guymer GP, McDonald WJ, Faith DP, Erickson D, Kress WJ. 2015. Mapping biodiversity and setting conservation priorities for SE Queensland's rainforests using DNA barcoding. PLoS One, 10:e0122164.

Shen XX, Liang D, Wen JZ, Zhang P. 2011. Multiple genome alignments facilitate development of NPCL markers: a case study of tetrapod phylogeny focusing on the position of turtles. Mol Biol Evol, 28:3237–3252.

Sievers F, Wilm A, Dineen D, Gibson TJ, Karplus K, Li W, Lopez R, McWilliam H, Remmert M, Soding J, et al. 2011. Fast, scalable generation of high-quality protein multiple sequence alignments using Clustal Omega. Mol Syst Biol, 7:539.

Smith MA, Fisher BL, Hebert PD. 2005. DNA barcoding for effective biodiversity assessment of a hyperdiverse arthropod group: the ants of Madagascar. Philos Trans R Soc Lond B Biol Sci, 360:1825–1834.

Sonet G, Jordaens K, Nagy ZT, Breman FC, De Meyer M, Backeljau T, Virgilio M. 2013. Adhoc: an R package to calculate ad hoc distance thresholds for DNA barcoding identification. Zookeys:329–336.

Spasojevic T, Kropf C, Nentwig W, Lasut L. 2016. Combining morphology, DNA sequences, and morphometrics: revising closely related species in the orb-weaving spider genus Araniella (Araneae, Araneidae). Zootaxa, 4111:448–470.

Stamatakis A. 2014. RAxML version 8: a tool for phylogenetic analysis and post-analysis of large phylogenies. Bioinformatics, 30:1312–1313.

Sutou M, Kato T, Ito M. 2011. Recent discoveries of armyworms in Japan and their species identification using DNA barcoding. Mol Ecol Resour, 11:992–1001.

Tanzler R, Sagata K, Surbakti S, Balke M, Riedel A. 2012. DNA barcoding for community ecology—how to tackle a hyperdiverse, mostly undescribed Melanesian fauna. PLoS One, 7:e28832.

van Velzen R, Weitschek E, Felici G, Bakker FT. 2012. DNA barcoding of recently diverged species: relative performance of matching methods. PLoS One, 7:e30490.

Vences M, Thomas M, Bonett RM, Vieites DR. 2005. Deciphering amphibian diversity through DNA barcoding: chances and challenges. Philos Trans R Soc Lond B Biol Sci, 360:1859–1868.

Virgilio M, Jordaens K, Breman FC, Backeljau T, De Meyer M. 2012. Identifying insects with incomplete DNA barcode libraries, African fruit flies (Diptera: Tephritidae) as a test case. PLoS One, 7:e31581.

Ward RD, Zemlak TS, Innes BH, Last PR, Hebert PD. 2005. DNA barcoding Australia's fish species. Philos Trans R Soc Lond B Biol Sci, 360:1847–1857.

Witt JD, Threloff DL, Hebert PD. 2006. DNA barcoding reveals extraordinary cryptic diversity in an amphipod genus: implications for desert spring conservation. Molecular ecology, 15:3073–3082.

Yang Z, Rannala B. 2010. Bayesian species delimitation using multilocus sequence data. Proceedings of the National Academy of Sciences of the United States of America, 107:9264–9269.

Young AD, Lemmon AR, Skevington JH, Mengual X, Stahls G, Reemer M, Jordaens K, Kelso S, Lemmon EM, Hauser M, et al. 2016. Anchored enrichment dataset for true flies (order Diptera) reveals insights into the phylogeny of flower flies (family Syrphidae). BMC Evol Biol, 16:143.

Yuan H, Jiang J, Jimenez FA, Hoberg EP, Cook JA, Galbreath KE, Li C. 2016. Target gene enrichment in the cyclophyllidean cestodes, the most diverse group of tapeworms. Mol Ecol Resour.

Zhao J, Li C, Zhao L, Wang W, Cao Y. 2008. Mitochondrial diversity and phylogeography of the Chinese perch, Siniperca chuatsi (Perciformes: Sinipercidae). Mol. Phylogenet. Evol., 49:399–404.

Zhou C, Yang Q, Cai D. 1988. On the classification and distribution of the sinipercinae fishes (family Serranidae). Zoological Research, 9:113–125.

